# What drives interaction strengths in complex food webs? A test with feeding rates of a generalist stream predator

**DOI:** 10.1101/259697

**Authors:** Daniel L. Preston, Jeremy S. Henderson, Landon P. Falke, Leah M. Segui, Tamara J. Layden, Mark Novak

## Abstract

Describing the mechanisms that drive variation in species interaction strengths is central to understanding, predicting, and managing community dynamics. Multiple factors have been linked to trophic interaction strength variation, including species densities, species traits, and abiotic factors. Yet most empirical tests of the relative roles of multiple mechanisms that drive variation have been limited to simplified experiments that may diverge from the dynamics of natural food webs. Here, we used a field-based observational approach to quantify the roles of prey density, predator density, predator-prey body-mass ratios, prey identity, and abiotic factors in driving variation in feeding rates of reticulate sculpin (*Cottus perplexus*). We combined data on over 6,000 predator-prey observations with prey identification time functions to estimate 289 prey-specific feeding rates at nine stream sites in Oregon. Feeding rates on 57 prey types showed an approximately log-normal distribution, with few strong and many weak interactions. Model selection indicated that prey density, followed by prey identity, were the two most important predictors of prey-specific sculpin feeding rates. Feeding rates showed a positive, accelerating relationship with prey density that was inconsistent with predator saturation predicted by current functional response models. Feeding rates also exhibited four orders-of-magnitude in variation across prey taxonomic orders, with the lowest feeding rates observed on prey with significant anti-predator defenses. Body-mass ratios were the third most important predictor variable, showing a hump-shaped relationship with the highest feeding rates at intermediate ratios. Sculpin density was negatively correlated with feeding rates, consistent with the presence of intraspecific predator interference. Our results highlight how multiple co-occurring drivers shape trophic interactions in nature and underscore ways in which simplified experiments or reliance on scaling laws alone may lead to biased inferences about the structure and dynamics of species-rich food webs.

## Introduction

Species interaction strengths typically vary over orders of magnitude, with most food webs containing few strong and many weak interactions (Wootton and Emmerson 2005). Such variation has important consequences for basic and applied ecological questions. For instance, the distribution and magnitudes of interaction strengths are important in predicting indirect net effects and the consequences of perturbations in food webs (Montoya et al. 2009, Novak et al. 2016), resolving relationships between network complexity and stability (Allesina and Tang 2012, Gellner and McCann 2016), and informing the management and conservation of interacting populations (Soulé et al. 2005). Quantifying the multiple factors that generate and maintain variation in the strength of species interactions is therefore central to understanding, predicting, and managing community dynamics.

A suite of factors has been advanced as drivers of interaction strength variation in food webs. Foremost is the density of interacting species, often formalized using predator functional response models (Jeschke et al. 2002). Functional response models generally predict that predator feeding rates increase with prey density and decrease with conspecific predator density, although the specific forms of the functions and the relative roles of prey vs. predator densities have been debated (Abrams and Ginzburg 2000). The characterization of predator feeding rates with functional response models represents a flexible framework for conceptualizing and quantifying trophic interaction strengths (Murdoch and Oaten 1975, Kéfi et al. 2012). Despite their utility, however, a considerable amount of unexplained variation in predator feeding rates, and interaction strengths defined more broadly, often remains after accounting for species densities.

Taxonomic identity and species traits, especially body size, have been invoked as additional drivers of variation in feeding rates. Predator size, prey size, and/or predator-prey size ratios, have been linked to variation in feeding rates directly, or to attack rate and handling time parameters within functional response models (Emmerson and Raffaelli 2004, Vucic-Pestic et al. 2010, Schmitz and Price 2011, Kalinkat et al. 2013). For example, allometric scaling, where biological rates vary with body size via power laws, has become a common approach for inferring interaction strengths in the absence of direct measures (e.g., Berlow et al. 2009). In addition to metabolic scaling relationships, there has been empirical support for hump-shaped relationships between predator-prey body mass ratios and feeding rates (Brose et al. 2008, Brose 2010). Lastly, a range of other characteristics beyond body size, including behavioral, chemical, or morphological traits that minimize predation risk (i.e. anti-predator defenses) can also influence trophic interaction strengths (Klecka and Boukal 2013, Kalinoski and DeLong 2016). Because such traits are often conserved (or correlated) across related species, the taxonomic identity of the prey or predator can drive significant variation in interaction strength (Rall et al. 2011).

While much focus has remained on characteristics of the interacting species themselves, the environment in which interactions occur may exert equally large controls on interaction strength. Many environmental factors, including light levels, habitat complexity, and especially temperature, have been shown as important drivers of trophic interaction strength (Pawar et al. 2012, Gilbert et al. 2014, Byers et al. 2017). For instance, water clarity has strong effects on the feeding rates of fish on zooplankton (Wissel et al. 2003); vegetation cover mediates rates of predation by wolves on elk (Kauffman et al. 2007); and moonlight affects kill rates of foxes on hares (Griffin et al. 2005). Taken together, these studies highlight the importance of characterizing variation in species interaction strengths across space and time (Chamberlain et al. 2014).

Although it is becoming widely recognized that interaction strengths are the outcome of dynamic biotic and abiotic factors, empirical tests of the relative roles of multiple drivers remain rare, particularly within the context of complex food webs (Wood et al. 2010). In large part, this disparity stems from challenges associated with measuring interaction strengths in nature. Limitations of prior work include a focus on one or two hypothesized drivers of interaction strength variation in isolation, and perhaps most importantly, the reliance on experiments entailing oversimplified community structures (e.g., low prey diversity) and unnatural environmental conditions (e.g., laboratory and mesocosm trials)(Carpenter et al. 1996). For instance, factors associated with study design, such as experimental duration (Li et al. in press) and the size of the feeding arena (Uiterwaal et al. 2017) have had strong effects on observed predator feeding rates. The large number of interactions occurring in complex food webs also makes experiments that manipulate the abundance or presence of species intractable. These challenges create disconnects between existing theory, results of laboratory studies, and the dynamics of real food webs (Wootton and Emmerson 2005, Novak and Wootton 2008).

In the present study we used an observational approach to quantify the feeding rates of a generalist predatory sculpin (*Cottus perplexus*) on its diverse suite of prey in the natural context of their species-rich stream food webs. We used model selection to compare the relative importance of five factors in driving *in situ* variation in predator feeding rates – predator and prey densities, predator-prey body mass ratios, prey taxonomic identity, and variation in the abiotic environment. We hypothesized that increases in prey density, and decreases in predator density, would be associated with increased prey-specific feeding rates. We further predicted that feeding rates would show a hump-shaped relationship with predator-prey body mass ratios, while prey taxonomic identity and abiotic factors would help explain additional variation beyond density and body size. We found that a full model including all five variables best explained prey-specific feeding rates, with prey density and taxonomic identity being the two most important factors. Our results emphasize the difficulty of extrapolating from lab-based studies of feeding rates to the field and underscore the need to empirically validate functional response models over realistic gradients of predator-prey characteristics and environmental factors.

## Methods

### Study System

Reticulate sculpin (*Cottus perplexus*) are benthic, generalist stream predators that typically reach an adult size of ~80 mm (Bond 1963, Pasch and Lyford Jr 1972, Finger 1982). We quantified feeding rates of reticulate sculpin at nine sites in Berry, Oak and Soap Creeks within Oregon State University’s McDonald-Dunn Research Forest northwest of Corvallis, Oregon (Fig. 1). Our study streams were ~1 to 3 m in width and flow through mixed coniferous forest. These streams support a diverse fauna of more than 325 benthic macroinvertebrate species (Anderson and Hansen 1987), as well as cutthroat trout (*Oncorhyncus clarkii*), Pacific giant salamanders (*Dicamptodon tenebrosus*), western brook lamprey (*Lampetra richardsoni*), and signal crayfish (*Pacifastacus leniusculus*).

**Figure 1.**
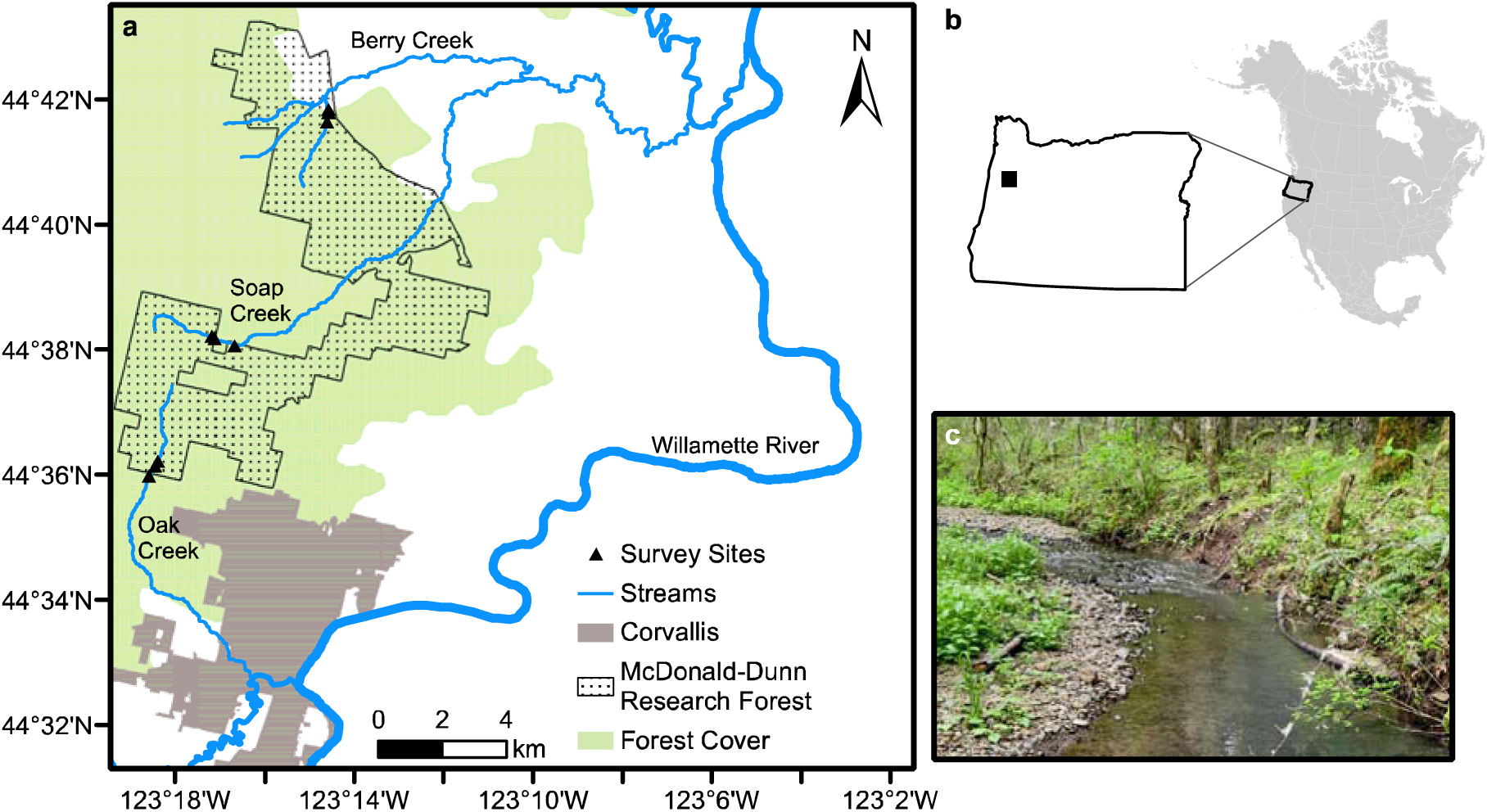
Map showing nine field sites at three streams where sculpin diets were surveyed (a). The map shows Oak Creek, Berry Creek and Soap Creek, which originate in the McDonald-Dunn Research Forest and flow into the Willamette River. The upper right inset shows the location of the study area in western Oregon, USA (b) and the lower right inset shows one of the study reaches at Berry Creek (c).

### Estimating Feeding Rates

Our approach for estimating sculpin feeding rates combines predator gut contents surveys with estimates of prey identification times (i.e. the time period over which prey remain identifiable in the stomach of a predator individual). Prey identification times allow diet counts to be converted into prey-specific feeding rates (i.e., prey consumed predator^−1^ time^−1^). Our approach represents a generalization of that described in Novak et al. 2017 (which itself follows from Novak and Wootton 2008 and Wolf et al. 2017) from predators that feed on a single prey item per observable feeding event to predators that feed on multiple prey items per observable feeding event (i.e. predator stomachs containing multiple prey items) (see also Woodward et al. 2005a). Prey-specific sculpin feeding rates were estimated for each site as

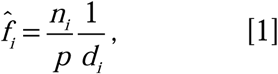

where 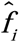 is the population-level mean feeding rate, *n_i_* is the number of prey items of species *i* found in a sample of *p* predator stomachs, and *d_i_* is prey *i*’s estimated identification time in the gut of the predator. With this approach, feeding rate estimates are independent of prey densities, and are dependent on body sizes only through their effects on prey identification times (see below).

### Field Surveys

In midsummer, we surveyed three reaches each at Berry, Oak and Soap Creeks (nine total sites; Fig 1) that measured ~45 m in length and contained a combination of riffle and pool habitat. At each site we quantified a suite of abiotic variables, including stream discharge, canopy cover, substrate size, water temperature, and stream width (see Supplemental Materials for details). We also collected ten Surber samples (0.093 m^2^ in area each) spaced evenly along each reach to quantify densities of benthic macroinvertebrate prey, which we preserved in 70% ethanol on-site and later identified in the laboratory (most to family, using Merritt et al. 2008).

To obtain diet information (*n_i_* and *p* in eqn. 1), a crew of four researchers used a backpack electroshocker (Smith-Root LR20B), a block net (1.0 × 1.0 m) and two dip nets (0.30 × 0.25 m) to systematically collect reticulate sculpin from each reach. Each captured sculpin was anesthetized, weighed, measured for total length, lavaged nonlethally to obtain stomach contents (using a 60 cc syringe with a blunt 18 gauge needle), and then released after a recovery period in aerated stream water. Sculpin stomach contents were preserved in 70% ethanol, and later identified and measured for total body length (see Table S1 for taxonomic resolution). Because partial digestion prevented every identifiable prey item from being measured, we applied prey-specific mean values – from either the relevant site when possible, or the entire dataset – to unmeasured prey items. We determined length-to-dry mass regressions or obtained these from published sources to estimate the dry mass of prey (Table S2) and converted sculpin wet mass into dry mass using a conversion factor of 0.24 (Lantry and O’Gorman 2007). To estimate each site’s sculpin density, we adjusted the catch totals from electroshock surveys by habitat-specific (pool or riffle) catch efficiencies estimated from mark-recapture surveys performed in each stream (see Supplemental Materials).

### Prey Identification Times

We estimated the prey identification times (*d_i_* in eqn. 1) on the basis of laboratory feeding trials during which a total of 356 sculpin were fed 879 prey items and then lavaged over time to determine whether prey remained identifiable or not. Detailed methods and analyses for determining prey identification times are provided in Preston et al. 2017 (see also Supplemental Information for a summary). Prey identification times were estimated as taxon-specific functions of water temperature, predator body size, and prey body size. We fed sculpin prey items from ten prey types, including mayflies (Ephemeroptera), stoneflies (Plecoptera), caddisflies (Trichoptera), flies (Diptera), beetles (Coleoptera), worms (Annelida), crayfish (*Pacifascatus leniusculus*), conspecific sculpin, conspecific sculpin eggs, and snails (*Juga plicifera*) (Table S3). We fit Weibull survival curves to observed prey statuses (identifiable or not) as a function of the three covariates (Klein and Moeschberger 2005). Coefficients from these functions were then used with covariate information from our field surveys to estimate prey identification times for each prey item based on the mean of the corresponding probability density function (Table S4, Fig. S1). For prey types that were not used in laboratory trials, we used survival function coefficients from a morphologically similar type of prey (e.g., for megalopterans we used coefficients estimated for trichopterans; see Table S1 and Supplemental Materials for additional details).

### Analyses

The primary aim of our analysis was to determine the relative importance of prey density, predator density, predator-prey body mass ratios, prey identity (i.e., taxonomic order), and abiotic environmental variables in driving variation in prey-specific sculpin feeding rates. The response variable in all analyses was the mean prey-specific feeding rates at each stream site. Our overall approach was to compare the relative fits of a full model with all five covariates, models with each covariate removed, and an intercept only null model (seven total models). We assessed the relative effect of dropping each variable using Akaike Information Criterion (AIC), Generalized Cross Validation scores (GCV), and adjusted r-square values (Burnham and Anderson 2002, Zuur 2009).

To determine the relationship between the five predictor variables and sculpin feeding rates, we used generalized additive mixed models (GAMMs). We included a random intercept term for reach nested within stream to account for the hierarchical structure of our survey design (Zuur 2009). For prey identity, prey items were grouped into 15 taxonomic orders (Table S1). To assess whether the sample size within orders affected the results, we also analyzed the same dataset after omitting seven orders with less than five feeding rate estimates each (Supplemental Materials). We incorporated body sizes using mean values at the site level based on individual measurements of each predator and their prey items. For prey items that were not detected in Surber samples (including terrestrial taxa; Table S1), we were unable to estimate prey densities and therefore omitted corresponding feeding rates from analyses (60 of 289 total feeding rates). To incorporate the abiotic variables into our analysis, we conducted a principal components analysis with site-level mean values for stream discharge, canopy cover, substrate size, water temperature, and stream width, and used the first principal component as a linear predictor variable in the GAMMs. We included a smoothing term for prey density and for predator/prey body mass ratios (using cubic regression splines) because we hypothesized that both variables could be linked to feeding rates in a nonlinear relationship. To determine the optimal amount of smoothing, we used generalized cross validation with the *mgcv* package in R (Wood 2017). We log-transformed feeding rates, prey densities, predator densities, and body mass ratios in all analyses to improve model assumptions. Correlation plots of predictor variables and plots of model residuals are provided in the Supplemental Materials (Figs. S2 and S3).

## Results

### Sculpin Stomach Contents

We lavaged a total of 778 reticulate sculpin and found 6988 identifiable prey items. Of the sculpin with identifiable prey, the mean number of prey items per fish was 9.5, with a range from one to 318 (median = 7, SD = 15.5). Forty sculpin (5.1%) did not contain identifiable prey. Across all sites we observed 57 prey types, most identified to the family level (Table S1). Within a given site, the total number of prey types observed ranged from 26 to 38 (see Fig. S4 for species accumulation curves). The majority of the individual prey items (94.6%), and the majority of prey types (68.4%), belonged to five orders (Coleoptera, Diptera, Ephemeroptera, Plecoptera and Trichoptera).

### Field Survey Covariates

Of the 57 observed prey types, 44 were found in surber samples at one or more sites, thereby allowing the estimation of prey densities (Table S1). Prey densities varied by three orders of magnitude, from 1 to > 1.1 × 10^3^ individuals m^−2^ (mean = 77, median = 24, SD = 150). Predator-prey body mass ratios varied by four orders of magnitude, from 1.9 × 10^1^ to 3.6 × 10^5^ (mean = 2.1 × 10^4^, median = 4.9 × 10^3^, SD = 4.9 × 10^4^). The distributions of prey densities and predator-prey body mass ratios were both highly skewed towards smaller values (Fig. 2). The estimated densities of sculpin ranged from 2.0 to 3.9 individuals m^−2^ (mean = 2.74, SD = 0.67) across sites. Variation in abiotic variables across sites was driven primarily by stream discharge, canopy cover, and stream width (Fig. S5). These three variables corresponded most closely to the first principal component of the PCA analysis, which explained 40% of the variation in the environmental data.

**Figure 2.**
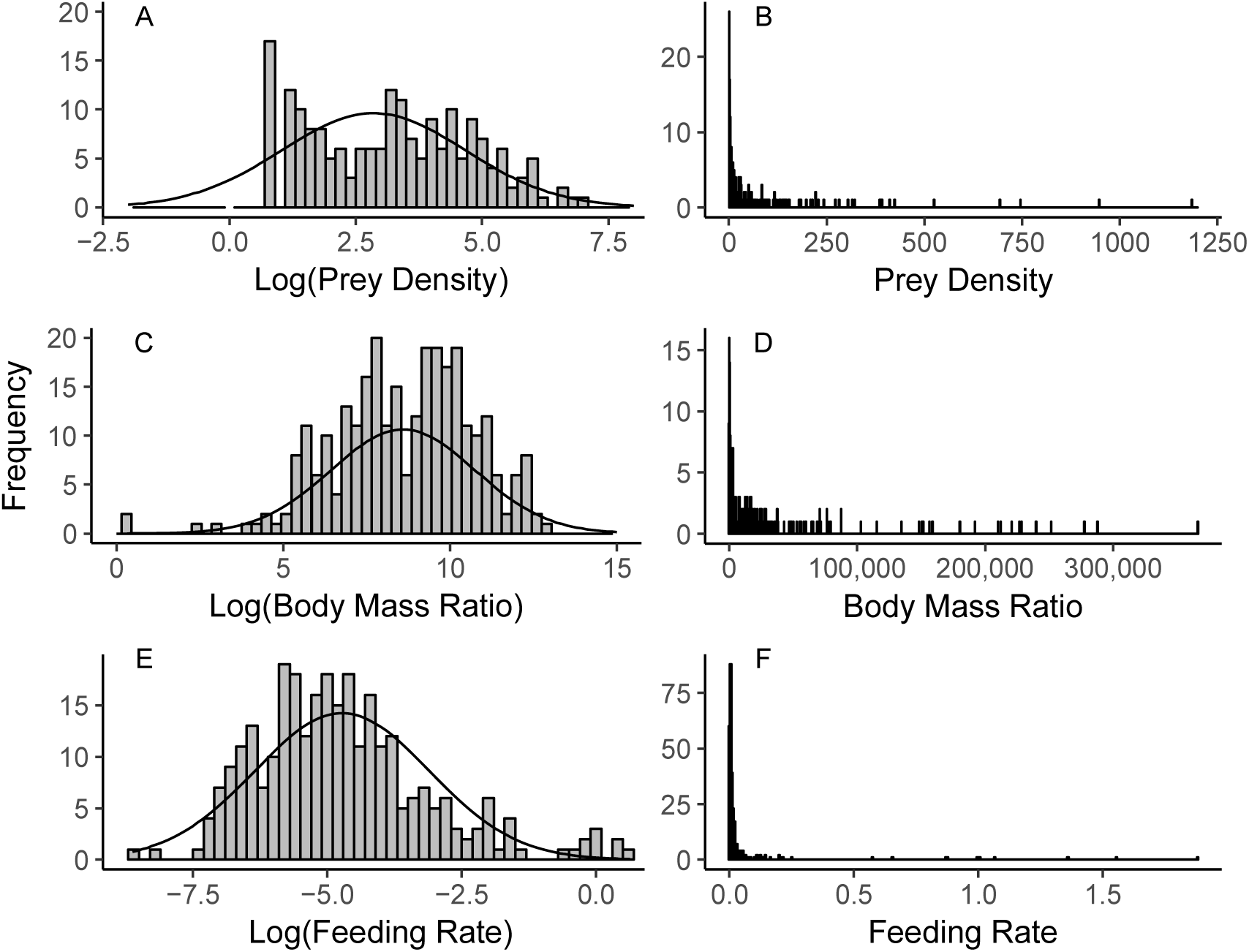
Distributions of log-transformed prey densities (a), untransformed prey densities (b), log-transformed predator-prey body mass ratios (c), untransformed body mass ratios (d), log-transformed sculpin feeding rates (e), and untransformed feeding rates (f). A normal curve parameterized with the observed mean and standard deviation from the empirical data is overlaid on the transformed data.

### Feeding Rates

Combining laboratory data on prey identification times with field diet information resulted in estimates of feeding rates for 289 prey type-by-site combinations. The prey identification times varied considerably across prey types, ranging from ~1 hr for annelid worms to ~60 hrs for snails (Fig. 3, Fig. S6). The wide range in prey identification times contributed to variation in the estimated feeding rates. For instance, annelid worms and signal crayfish occurred with similar frequencies in the sculpin stomach contents, with a mean number of 0.023 (worms) and 0.022 (crayfish) per sculpin, but differed by ~29 hrs in their mean prey identification times and thereby had a 50-fold difference in estimated mean feeding rates (worms = 3.3 × 10–^2^; crayfish = 7.2 × 10–^4^ prey consumed sculpin^−1^ hour^−1^).

**Figure 3.**
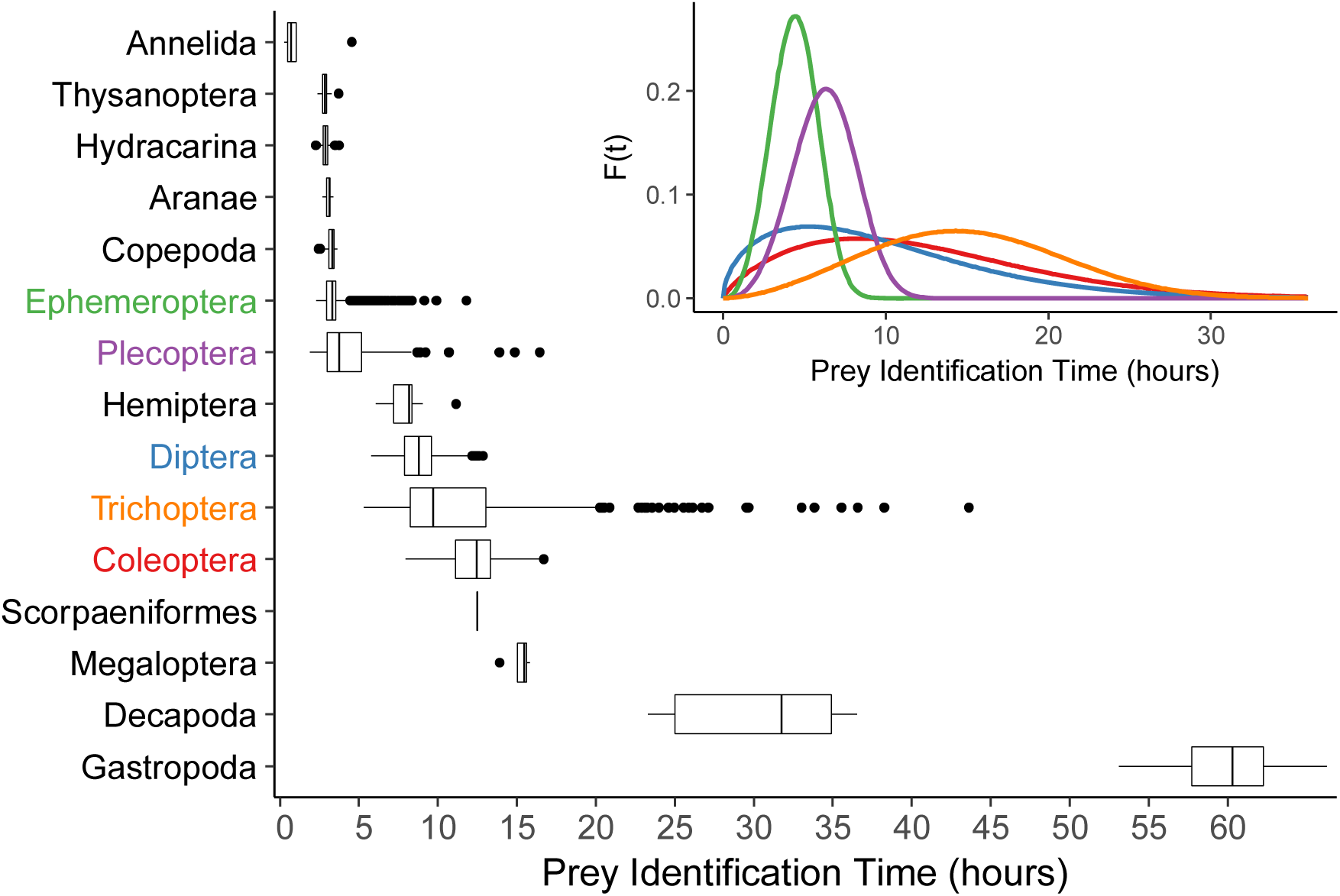
Boxplot showing estimated prey identification times for the 15 prey orders observed in sculpin stomach contents. The prey identification times in the boxplot are calculated from the mean of Weibull probability density functions for each prey item, which are shown in the inset panel for the five most common prey orders (colors correspond to taxa on the y-axis of the boxplot; see Supplemental Materials for density functions for other prey taxa). The functions in the inset were parameterized based on laboratory feeding trials in combination with mean covariate values from the field data. On the boxplot, the upper and lower hinges correspond to the first and third quartiles, the horizontal line is the median, the whiskers extend to the highest and lowest values within 1.5 times the interquartile range, and outliers are shown as solid points.

The distribution of feeding rates was approximately log-normal, with many low and a few relatively high values (Fig. 2e, 2f). That said, we observed more feeding rates than predicted for a log-normal on the left side of the distribution and a cluster of very high feeding rates at the far right side of the distribution. Feeding rates varied over four orders of magnitude, from 1.8 × 10–^4^ (freshwater bivalves) to 1.88 (baetid mayflies), with a mean of 3.5 × 10–^2^ (median = 6.7 × 10–^3^, S.D = 0.15). Body mass ratios and prey densities also showed highly skewed distributions (Fig. 2).

The full model predicting sculpin feeding rates with all five covariates fit the data better than, or equally, to all the other models (AIC = 656.1, r^2^ = 0.62; Table 1). Of the covariates, dropping prey density from the full model resulted in the largest decrease in relative mode fit (ΔAIC = 100.1). The next most important covariate was prey identity (ΔAIC = 51.1 when dropped from the full model). Dropping each of the other three covariates resulted in smaller changes in model fit (ΔAIC = 7.3 for predator-prey body mass ratios; ΔAIC = 1.1 for predator density; ΔAIC = -0.5 for PC1 of the abiotic factors). The intercept only null model performed the most poorly (ΔAIC = 185.5 relative to the full model). A smaller dataset omitting prey orders with low sample sizes did not alter the main results (Supplemental Materials and Table S5).

**Table 1.** Model comparisons indicating the relative importance of five hypothesized drivers of variation in sculpin feeding rates. The models include a full model with all five variables, five models dropping each of the five predictors separately, and an intercept only null model. The table includes the log likelihood values, Akaike Information Criterion (AIC), the change in AIC from the full model (ΔAIC), generalized cross validation scores (GCV), and an adjusted R^2^ statistic. All of the models include smoothing terms for prey density and body mass ratios, and a random intercept term for survey site nested within stream.

| Model | Log Likelihood | AIC | ΔAIC | GCV | Adjusted $R^2$ |
| --- | --- | --- | --- | --- | --- |
| Full Model | -300.0 | 656.1 | 0.0 | 1.180 | 0.617 |
| Full Model Without PC1 | -300.8 | 655.6 | -0.5 | 1.176 | 0.616 |
| Full Model Without Predator Density | -299.0 | 657.2 | 1.1 | 1.188 | 0.617 |
| Full Model Without Mass Ratios | -307.7 | 663.4 | 7.3 | 1.214 | 0.598 |
| Full Model Without Prey Order | -334.6 | 707.2 | 51.1 | 1.476 | 0.498 |
| Full Model Without Prey Density | -353.7 | 756.2 | 100.1 | 1.855 | 0.386 |
| Intercept Only Model | -417.1 | 841.6 | 185.5 | 2.708 | 0.010 |

Of the five hypothesized drivers of feeding rate variation, prey density was most closely correlated with observed feeding rates, showing an accelerating relationship (Fig. 4a). Over the range of prey densities observed, the predicted sculpin feeding rate increased by ~85 fold. The two prey taxa with the highest densities at each site – baetid mayflies and midge larvae – corresponded with the highest sculpin feeding rates. Feeding rates demonstrated a unimodal relationship with predator-prey body mass ratios, peaking at values around 2.2 × 10^4^ (Fig. 4b). Variation in feeding rates was highest at intermediate body mass ratios. A small number of very large prey items (signal crayfish and conspecific sculpin) resulted in a wide confidence interval for predicted feeding rates at the smallest observed predator-prey body mass ratios (Fig. 4b). Predator density was negatively correlated with sculpin feeding rates, with a doubling in density from two to four sculpin m^−2^, associated with a 2.5-fold decrease in mean feeding rates (Fig. 4c). There was also considerable variation in feeding rates associated with prey identity, ranging from a mean rate of 0.33 for mayflies to 0.007 for crayfish (Fig. 4e). Lastly, feeding rates did not vary between sites in a manner that was correlated with first principal component of the abiotic factors (Fig. 4d).

**Figure 4.**
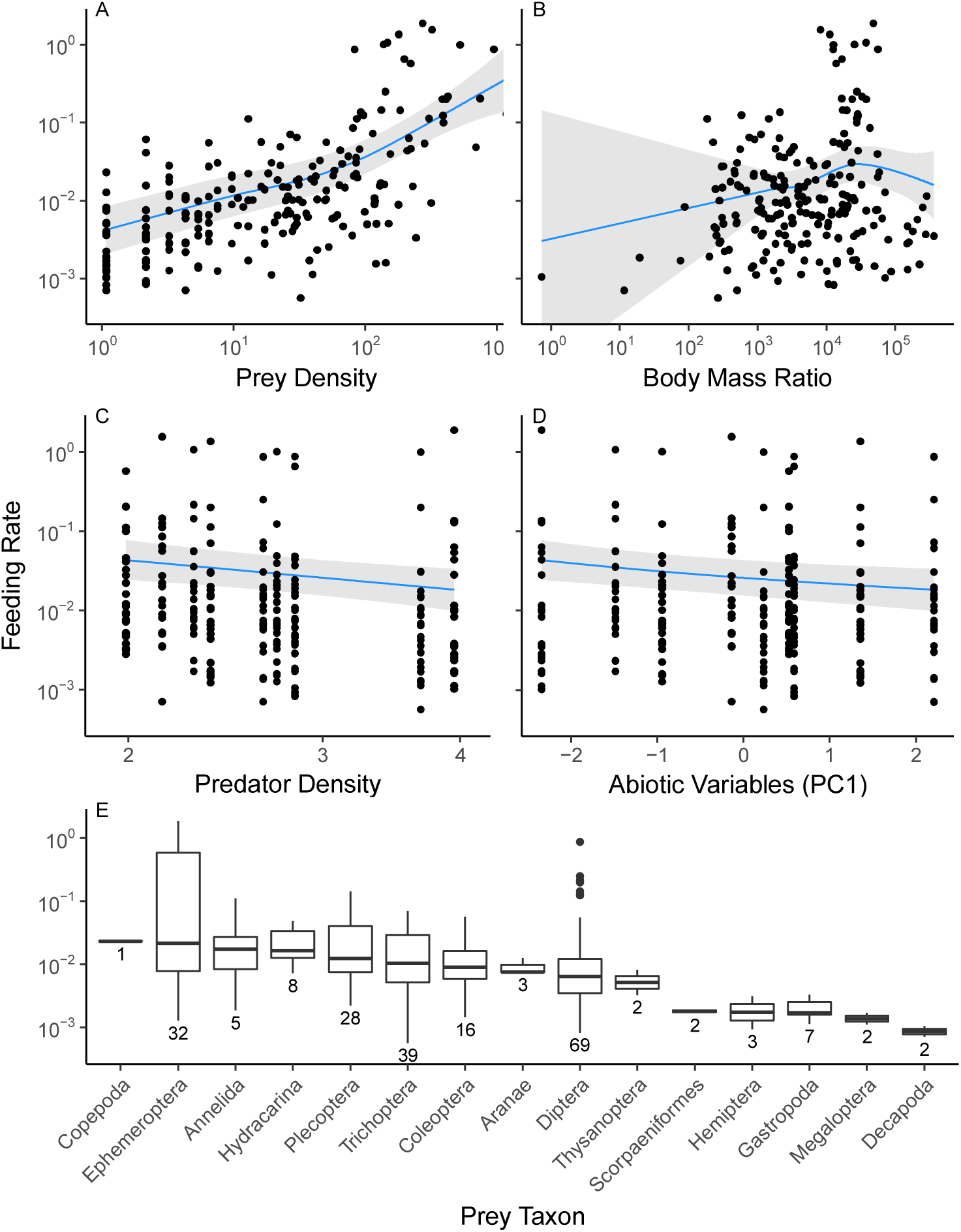
Five hypothesized drivers of variation in sculpin feeding rates. The y-axis for all panels is the prey-specific feeding rates at nine stream sites. The variables on each plot include prey density (prey individuals m^−2^) (a), predator-prey body mass ratios (b), predator density (sculpin m^−2^) (c), the first principal component from a PCA of five abiotic stream variables (d), and prey taxonomic order (e). Note the y-axis for all panels, and the x-axis for panels a,b, and c, are on a log scale. The regression lines correspond to a generalized additive mixed model including all five predictor variables (the ‘full model’ in Table 1, using means of continuous variables and Plecoptera as a representative prey order for predicted values). The grey bands correspond to 95% confidence intervals. For the boxplot, the number of taxa within each prey order is indicated below the boxes, with the groups being ordered by their median values. The upper and lower hinges correspond to the first and third quartiles, the horizontal line is the median, the whiskers extend to the highest and lowest values within 1.5 times the interquartile range, and outliers are shown as solid points.

## Discussion

A wide range of factors – including characteristics of predators, their prey, and the environment – have been linked to variation in the strength of trophic interactions. As a result, a growing suite of functional response models that differentially incorporate these factors has been advanced to characterize predator feeding rates. To date, however, most efforts to understand the relative roles of multiple factors in driving variation in feeding rates have relied on simplified experiments that are far removed from the complexity of natural food webs. By quantifying feeding rates of a focal generalist predator *in situ*, we assessed the relative importance of five primary factors that have been predicted to drive variation in trophic interaction strengths under natural conditions. Our results provide support for several predictions of predator-prey theory, but also emphasize key areas in need of further study to strengthen linkages between functional response models and the dynamics of trophic interactions in the field.

Consistent with prior studies of interactions strengths, we observed a skewed distribution of feeding rates, with few strong and many weak interactions. This pattern has been observed whether interaction strength measures are based on energy flow (e.g, Cross et al. 2013, Bellmore et al. 2015), experimental indices of interaction strength (e.g., Wood et al. 2010), or feeding rates (e.g., Wootton 1997). Recent theory suggests that as food webs increase in complexity, the skewed distribution of interaction strengths should become even more pronounced (Iles and Novak 2016). Potential drivers of interaction strengths often show skewed distributions as well, including species abundances and body sizes (Kozlowski and Gawelczyk 2002, McGill et al. 2007). Feeding rates, prey densities, and body-mass ratios all had highly skewed distributions in our study. While the consequences of skewed interaction strength distributions for community dynamics have received ample attention, the mechanisms driving distribution shape remain less well understood. Statistical processes may play a role, as power law and log-normal distributions can result from the multiplicative processes that underlie feeding rates (Limpert et al. 2001). It has also been posited that the skewed distribution could result from a tendency for communities to move towards stable configurations over time (e.g., via colonization, extinction, and/or evolution)(Wootton and Emmerson 2005, Borrelli et al. 2015). Our results add to this discussion by testing the relative roles of multiple biological factors in driving variation in *in situ* trophic interaction strengths.

Among the five predictor variables, prey density was most strongly correlated with variation in sculpin feeding rates. This result was consistent with the positive association between encounter rate and prey density predicted by all functional response models at low relative prey densities. At high prey densities, however, the most commonly applied functional response models (e.g., Holling type II and III forms) predict that predator feeding rates should eventually saturate because feeding rates become limited by prey handling time rather than encounter rate (Holling 1959). We observed an increasing positive slope at the highest observed prey densities, which is opposite to the decreasing slope that would be consistent with predator saturation. This finding is likely the result of several factors. First, the time needed for sculpin to physically consume a typical prey item is very short (based on extensive laboratory observations); feeding rates are more likely to be limited by digestion processes and stomach capacity. Our field data, however, indicate that sculpin were well below their maximum stomach capacity. The highest number of observed prey items in one sculpin was 318 mayflies, with most sculpin having far fewer prey (mean of ~10). Taken together, these observations suggest that sculpin feeding rates in the field are rarely, if ever, limited by handling or digestion times, making saturating functional responses potentially unrealistic.

While predator feeding rates must theoretically become limited at some point by handling or digestion processes, the generality of saturating feeding rates often implied in the literature deserves further study within the context of realistic prey densities in natural food webs. Empirical support for saturating functional responses is based mostly on laboratory studies that may exceed the range of prey densities that occur in nature and involve simplified environments that maximize predator-prey encounter rates. While the functional form of a predator’s feeding rates will depend on the biology of the system, the specifics of our study system – i.e., short handling times, relatively small prey, and large stomach capacity – are not uncommon in other predator-prey systems (Jeschke et al. 2002). Even predators with long handling times may still rarely experience saturation. For instance, feeding rates of whelks in the marine intertidal, which exhibit classic Type II functional responses in laboratory experiments, have been shown to be better described by linear functional responses when quantified in the field (Novak 2010).

One possible factor contributing to the lack of predator saturation in this study system is the physical habitat in which trophic interactions occur. The complex three-dimensional matrix of the stream benthos, with a deep layer of cobble and gravel, may have resulted in much lower realized encounter rates than would a comparable prey density in a less complex habitat. Although prey densities reached > 1000 individuals m^−2^ for some taxa (Chironomidae), and exceeded 2800 invertebrate m^−2^ at some sites for all taxa combined, the actual number of prey coming into contact with a sculpin at a given point in time is probably much lower than these numbers reflect due to the spatial complexity of the habitat. Sculpin are known to move through the stream substrate at a considerable depth (Phillips and Claire 1966, Thomas 1973), consistent with our observations while conducting surveys. Physical complexity could therefore contribute to preventing saturation by lowering encounter rates at high prey densities. More broadly, recent work has indicated that the dimensionality (2D vs 3D) and physical complexity of the habitat can strongly mediate predator feeding rates in diverse systems (Pawar et al. 2012, Barrios-O’Neill et al. 2016).

Body sizes of predators and prey have often been invoked as a primary driver of trophic interaction strength (Woodward et al. 2005b, but see Wootton and Emmerson 2005). Including predator-prey body mass ratios slightly improved the relative fit of our model predicting sculpin feeding rates. Feeding rates were highest at body mass ratios of around 2.2 × 10^4^, and then decreased for smaller and larger prey. The relationship was not symmetrical, however, with the peak towards larger body mass ratios (i.e., smaller prey). Sculpin, like many predatory fishes, are gape limited and consume their prey whole, creating an upper size limit on prey that can be consumed (Tabor et al. 2007). This probably contributes to the higher feeding rates on relatively small prey. The hump-shaped relationship between feeding rates and body-mass ratios is consistent with foraging theory which posits that large prey are difficult to consume while very small prey are not energetically cost effective, resulting in the highest feeding rates at intermediate body-size ratios (Brose et al. 2008). We note, however, that there was a large amount of variation in feeding rates at intermediate prey body sizes. Such variation may not have been observed in past laboratory-based studies because predators were exposed to only a single prey species at a time.

Prey identity was the second most closely associated variable with variation in feeding rates, and exceeded the importance of body-mass ratios. Mean feeding rates varied by over four orders of magnitude across prey identities. We used taxonomic order as a grouping variable with the aim that it would represent conserved variation in prey traits within an order. While multiple prey traits (e.g., behavior, morphology) could underlie differences in prey-specific sculpin feeding rates, one key trait captured by this approach is anti-predator defenses in the form of external protection. Two of the three lowest observed feeding rates were on taxa that are relatively abundant but possess external protection – signal crayfish and *Juga* snails. These two taxa had the longest prey identification times in the laboratory (~30 and 60 hrs), suggesting that low digestibility may contribute to sculpin preferring other prey. In the laboratory, sculpin were reluctant to consume snails, often regurgitating them shortly after consumption (pers. obs.). In contrast, prey taxa with short identification times that were rapidly digested – including mayflies and annelid worms – had two of the three highest feeding rates and lack external protection. In general, we found that feeding rates were negatively associated with prey identification times (Fig. S8), which is an *a priori* expectation based on eqn. 1, but also a biologically reasonable prediction for generalist predators. Additionally, two of the four lowest feeding rates were on hemipterans and megalopterans, both possessing other forms of defenses that could possibly minimize predation by sculpin (toxins and formidable mandibles, respectively) (Merritt et al. 2008). The importance of prey traits in driving variation in predator feeding rates is beginning to be recognized and quantified in laboratory experiments (Rall et al. 2011, Klecka and Boukal 2013, Kalinoski and DeLong 2016, Uiterwaal et al. 2017). Our findings, coupled with prior work, suggest that prey traits beyond body mass are likely a fundamental driver of variation in predator feeding rates. Taxonomic- or traits-based approaches may therefore prove especially useful at explaining the large variation or divergences from predictions of metabolic theory often observed in relationships between body mass or temperature and feeding rates (Brose et al. 2008, Englund et al. 2011, Rall et al. 2012). The role of such traits will not be evidenced by experimental studies with one or a few prey types, underscoring the need to quantify interaction strengths in diverse communities.

Predator density had a relatively small effect on sculpin feeding rates, although the negative relationship was consistent with predator interference as encapsulated by most consumer-dependent functional response models (Skalski and Gilliam 2001). The density of conspecific predators can influence feeding rates through a variety of mechanisms, many involving changes in predator behavior (Abrams and Ginzburg 2000). This observation has fueled debate over whether and how functional responses should incorporate predator density, but few studies have assessed its importance in the field (Novak et al. 2017). Our results provide some support for the presence of predator interference in this system. Behavioral interactions between sculpin, and specifically a competitive hierarchy for feeding sites based on body size, have been observed in related stream sculpin species (Grossman et al. 2006). Sculpin are also territorial when guarding their eggs in the spring months, but this would have occurred prior to our surveys in this study (Bateman and Li 2001). There is also evidence that other predator species can influence sculpin feeding rates (Soluk 1993). Our stream sites support cutthroat trout and Pacific giant salamanders, both of which share invertebrate prey of sculpin (Falke et al. in prep). Future work on the roles of intra- and interspecific interactions with other predators in driving variation in feeding rates is needed to test the form and magnitude of predator dependence in this study system.

One consideration in the interpretation of our results is that we surveyed over a relatively small geographical area and timespan. As a result, our stream sites were fairly similar in abiotic factors, limiting our ability to detect differences in feeding rates driven by the environment. Surveys replicated over different times of year, with greater variation in temperature, stream flow, habitat type (pool vs. riffle), and cobble size would be more effective at detecting environmental effects. Furthermore, our focus on one predator taxon limits our ability to investigate effects caused by predator characteristics. For instance, datasets with a wide range of predator size across multiple taxa would be better suited for detecting relationships driven by consumer mass independent of prey mass (e.g., Barrios-O’neil et al. 2016). Nonetheless, the consistency of prey-specific feeding rates across sites in our study is notable. Also, the food webs we studied support a highly diverse prey base (>325 invertebrate species) and relatively few aquatic vertebrate predators (three species). These observations suggest that prey characteristics may drive more variation in community wide feeding rates than predator characteristics, although this remains to be tested. The high prey richness in our study system also contrasts with most empirical food webs that typically entail predators with much lower numbers of prey species per predator (e.g., Dunne et al. 2002), and makes our dataset especially powerful for detecting the roles of prey characteristics.

Predator-prey theory has often outpaced realistic empirical tests of model predictions. Predator functional responses, for instance, have been largely based on laboratory experiments and have seen relatively little validation in natural food webs. Artificial effects of the laboratory environment, including experimental duration (Li et al. in press), the size of the arena used for feeding trials (Uiterwaal et al. 2017), or unrealistic community structure (e.g., low prey richness) have potential to strongly influence predator feeding rates. Additionally, scaling laws based on temperature and body mass have been applied to infer interaction strengths in food webs, despite sometimes equivocal support for the generality of their predictions (Wooton and Emmerson 2005, Rall et al. 2012). Our results reinforce that the scale of inference – in space, time, and taxonomic diversity of predators and prey – will have important implications for the factors generating variation in trophic interactions strengths. While scaling laws and generic predictions from functional response models may be of use in predicting broad patterns of linkage strength or energy flow, the trophic dynamics within a given food web will often diverge from these expectations due to unique taxon-specific traits or other local factors. By quantifying species interaction strengths within the context of complex food webs, empirical studies have the potential to dissolve many more of the disconnects between theory, laboratory studies, and the dynamics of trophic interactions in nature.

## Acknowledgements

For assistance with data collection we thank Madeleine Barrett, Alicen Billings, Daniel Gradison, Kurt Ingeman, Dana Moore, Arren Padgett, Wendy Saepharn, Alex Scharfstein, Johnny Schwartz, Isaac Shepard, Samantha Sturman, Ernesto Vaca Jr, and Beatriz Werber. Richard Van Driesche and Michael Bogan provided input on identifying aquatic invertebrates, Alison Iles provided comments that improved the manuscript, and Sarah Hart assisted with creating the map of the study sites. We thank the Oregon State University College of Forestry for providing access to field sites in the McDonald-Dunn Research Forest and for GIS data, particularly Steve Fitzgerald and Brent Klumph. Stan Gregory provided valuable input on our field sites and past research. Funding was provided by the National Science Foundation Grant DEB-1353827 and Oregon State University.

